# Invariant Differential Expression Analysis Reveals Mechanism of Cancer Resistance to Cell Cycle Inhibitors

**DOI:** 10.1101/2021.02.17.431607

**Authors:** Amit Chatterjee, Sonalisa Pandey, Ravikanth Danda, R Ranjith Kumar, S Maheswari, Vikas Khetan, Pukhraj Rishi, S Ramaprabhu, Sailaja V Elchuri, Debashis Sahoo

## Abstract

Retinoblastoma (RB) is a good model to study drug resistance to cell-cycle inhibitors because it is driven by mutations in the core components of cell-cycle, i.e, Rb gene. However, there is limited gene expression dataset in RB which has major reproducibility issues. We have developed invariant differential expression analysis (iDEA) that improves the state of the art in differential expression analysis (DEA). iDEA uses strong Boolean implication relationships in a large diverse human dataset GSE119087 (n = 25,955) to filter the noisy differentially expressed genes (DEGs). iDEA was applied to RB datasets and a gene signature was computed that led to prediction and mechanism of drug sensitivity. The prediction was confirmed using drugs-sensitive/resistant RB cell-lines and mouse xenograft models using CDC25 inhibitor NSC663284. iDEA improved reproducibility of differential expression across diverse retina/RB cohorts and RB cell-lines with different drug sensitivity (Y79/Weri vs NCC). Pathway analysis revealed WNT/β-catenin involved in distinguishing drug sensitivity to CDC25 inhibitor NSC663284. NSC663284 inhibited tumour cell proliferation in mouse xenograft model containing Y79 cells indicating novel therapeutic option in RB. Invariant differentially expressed genes (iDEGs) are robustly associated with outcome in diverse cancer datasets and supports for a fundamental mechanism of drug resistance.

## Introduction

Differential expression analysis (DEA) is a standard approach to identify differentially expressed genes (DEGs) between normal and disease human tissue^1,2^. DEA has been widely used in analyzing large scale biomedical data^3-8^. Reproducibility of the results has been a major issue that affect progress in understanding biological processes^9,10^. A previous study establish that current methods can give reproducible false positive findings that are driven by genetic regulation of gene expression, yet are unrelated to the trait of interest^11^. These problems persist in previous improvements on DEA^12-15^. We developed a new computational approach that performs an unbiased differential expression analysis combined with strong Boolean implication relationships^16^. We call this invariant differential expression analysis (iDEA) that identifies invariant differentially expressed genes (iDEGs) because these genes maintain strong Boolean implication relationships in almost all tissues regardless of their disease state. This method is so simple and general that it can be applied in any gene expression differential analysis context and complimentary to all previous improvements. We applied this approach to Retinoblastoma (RB) to demonstrate a proof of principle (Fig 1A). We demonstrate how this approach accelerate discovery of drug targets and potential mechanism of action.

**Figure 1:**
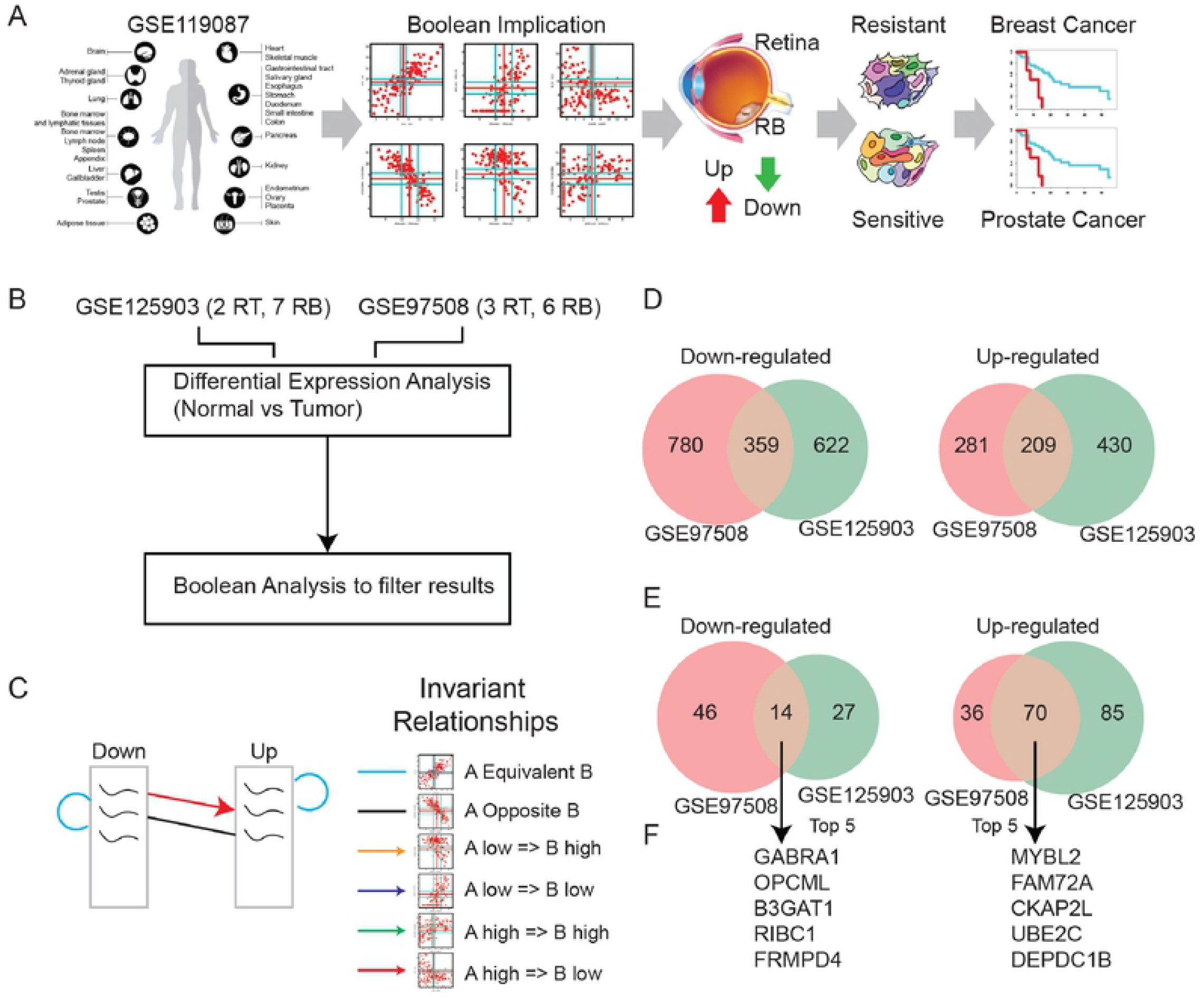
Study design and invariant differential expression analysis. (A) Study design overview that use a new computational approach to identify fundamental mechanism of cancer resistance in diverse cancer types. (B) Schematic approach to identify invariant differentially expressed genes between normal retina and retinoblastoma datasets: GSE125903 (2 control retina vs 7 retinoblastoma; RNASeq) and GSE97508 (3 control retina vs 6 retinoblastoma; microarray). (C) Schematic approach for filtering traditional differential analysis by using Boolean implication relationships (color coded) on a global diverse tissue dataset GSE119087 (n = 25,955; Affymetrix Human U133 Plus 2.0). The algorithm uses three criteria: (1) Boolean equivalent within each of the Down- and Up-regulated genes, (2) Boolean high => low from Down-regulated genes to up-regulated genes, (3) Boolean opposite from Down-regulated genes to up-regulated genes. (D) Overlap of differential expression analysis in two datasets: GSE125903 vs GSE97508. (E) Overlap of invariant differential expression analysis in two datasets: GSE125903 vs GSE97508. (F) Top five Down- and Up-regulated genes.

Retinoblastoma (RB) arises due to mutation in Retinoblastoma protein (Rb)^17^. Rb is a tumor suppressor protein and its mutation leads to pediatric intraocular cancer that originates from the neuroectodermal cells of the retina (RT)^18,19^. Rb tumor are caused by the inactivation of the pRb protein which is a tumor suppressor protein^20^. RB is classified into two types bilateral which is caused by a germline mutation of the RB1 gene and unilateral which is sporadic and caused by external factors such as mutagens, viruses like HPV^21^. Treatment of RB especially in high grade tumors involves enucleation followed by conventional adjuvant chemotherapy^22^. These chemotherapeutic agents target rapidly dividing cells including cancer cells and normal cells. Additionally, the use of chemotherapeutic agents has grave effects in the treatment of childhood cancers, which hinders the normal growth of effected child. Additionally, the RB disease progression leads to partial or complete loss of vision effecting the quality of life of these children^23^. Therefore, finding targeted therapy molecules in reducing RB tumour progression might reduce the side effects in these children. The therapeutic target identification used global transcriptomics and proteomic approaches^24-26^. The deregulated pathways identified in RB includes AKT signalling pathway, Wnt pathway, IGF signalling pathway and MAPK pathways^27^. Recently, RNA sequencing of RB tumours by us identified altered noncoding RNA and fusion transcripts. However, there is need for identifying additional therapy molecules for RB treatment^28^.

Phenotypic characterization of RB tumor cells has identified the presence of stem cell markers such as MDR1, ABCG2, Oct4, Nanog, ALDH1, and CD44^29^. Wnt signaling plays a pivotal role in the formation of RB tumor and can increase stem-like cells in RB tumors^30^. This pathway has been reported to be proto-oncogenic in most of the solid tumor whereas, this pathway is known to be tumor suppressor in RB^31,32^. iDEGs robustly classified retina vs retinoblastoma, severity in retinoblastoma, drug sensitivity in RB Cell lines (sensitive Y79/Weri^33,34^ vs resistant NCC RB51^35^). Pathway analysis of differentially expressed genes between these RB cell lines revealed WNT/β catenin to be an important player. WNT/β-catenin pathway has been shown previously to mediate drug resistance in tumors^36,37^. Therefore, it is essential to understand the WNT signalling pathway and its regulation by potential drug therapy molecules to identify novel therapy targets in RB. RB has been shown to up-regulate genes involved in cell cycle such as CDC25A/B/C^4,38-40^. Interestingly CDC25 has recently been identified as a target for diverse triple-negative breast cancers including *RB1/PTEN/P53*-deficient tumors^41^. However, small molecule CDC25 therapy for RB progression is not studied before. Additionally, the cross talk between WNT signaling and targeted therapy molecules that can increase RB cell apoptosis is not studied before. *In vitro* cell culture models like Y79, Weri and NCC RB51 are employed to understand the mechanism between WNT signaling and cell cycle progression inhibitors. We discovered that the RB cells lines express differential localization of transcription factors. The localization of β-catenin, a central molecule of WNT signaling and its interacting partners such as c-fos and c-jun was nuclear in Weri and Y79 RB cells. However, their localization is different in NCC cell line where the WNT signalling molecules were in the periphery of the cells. Furthermore, the WNT activated cell lines exhibited therapeutic response to the CDC25 small molecule inhibitor, NSC663284 while the NCC Rb51 could not respond to this therapy. Using WNT activated cell line Y79 we created the mouse xenograft model and used CDC25 small molecule inhibitor, NSC663284 as treatment for reducing tumor cell proliferation. Our results indicate for the first time that WNT signalling is needed for therapeutic response of CDC25 small molecule inhibitor. Finally, iDEGs were tested on diverse cancer datasets and association with outcome. iDEGs performed much better compared to DEGs (Differentially Expressed Genes) because of the invariant aspect of the analysis which helps translate the information from retina to other tissue types.

## Materials and Methods

### Sample collection

The present study was conducted at the Medical Research Foundation and Vision Research foundation Sankara Nethralaya, India. The study was approved by the Institutional Ethics Board. Ethics No. 247-2011-P. Normal adult retinas (n=3, age 30 yrs) were obtained from the enucleated eyeballs of cadaveric donors donated to C.U SHAH eye bank, Sankara Nethralaya. A part of the tumor from the enucleated eyeball was used for pathological evaluation and the remaining part was used for gene expression analysis by Q-PCR. The tumors were graded according to international retinoblastoma staging system for RB. The collected tumors were stored in liquid nitrogen until further use. Clinicopathological characteristics for 10 RB samples are given in supplementary table 1.

### Data Collection

Publicly available microarray and gene expression databases were downloaded from the National Center for Biotechnology Information (NCBI) Gene Expression Omnibus website (GEO) ^42-44^ and the European Molecular Biology Laboratory European Bioinformatics Institutes (EMBL-EBI) ArrayExpress website^45-47^. NCBI GEO and EMBL-EBI Array Express were searched for transcriptomic studies of retinoblastoma. If the dataset is not normalized, RMA (Robust Multichip Average)^48,49^ is used for microarrays and TPM (Transcripts Per Millions)^50,51^ is used for RNASeq data for normalization. We used log2(TPM+1) to compute the final log-reduced expression values for RNASeq data. Accession numbers for these crowdsourced datasets are provided (Supplemental Table 4). All of these datasets were processed using the Hegemon data analysis framework ^52-54^.

### Identification of Boolean implication relationships

Boolean implication analysis is performed in the large diverse human dataset GSE119087 (n = 25,955 human samples). First, gene expression levels were converted to Boolean values (high and low) using StepMiner algorithm (Fig. S1A)^55^. The expression values are sorted from low to high and a rising step function is fitted to the series to identify the threshold (Fig. S1A). Relationship between two genes was evaluated in the context of high and low values (Fig. S1B-G). These relationships are called Boolean implication relationships (BIRs) because they are represented by logical implication (=>) formula. BooleanNet statistics is used to assess the significance of the Boolean implication relationships^16^. S > 3 and p < 0.1 are the thresholds (False Discovery Rate < 0.0001) used on the BooleanNet statistics during the neuroblastoma data analysis to identify Boolean implication relationships. A noise margin of 2-fold change is applied around the threshold to determine intermediate values and these values are ignored during Boolean analysis.

### Differential Expression Analysis (DEA)

Differential expression analysis is performed using multiple t-tests in python scipy.stats.ttest_ind package (version 0.19.0) with Welch’s Two Sample t-test (unpaired, unequal variance (equal_var=False), and unequal sample size) parameters. Multiple hypothesis correction were performed by adjusting *p* values with statsmodels.stats.multitest.multipletests (fdr_bh: Benjamini/Hochberg principles). The results were independently validated with R statistical software (R version 3.6.1; 2019-07-05). Adjusted p-value threshold for GSE97508 was 0.1 and for GSE125903 was 0.15 (Fig. S1H).

### Invariant Differential Expression Analysis (iDEA)

Invariant Differential Expression Analysis (iDEA, Fig. S1I) is a new computational approach where standard differentially expressed genes are filtered using Boolean implication relationships on large diverse human dataset GSE119087 (n = 25,955 human samples). The Up/Down lists are first filtered by using Boolean Equivalent relationships. First, genes from list (Up/Down) that do not have Boolean Equivalent relationships with other genes from the same list (Up/Down) is removed. Second, up-regulated genes that do not have Boolean high => low or Opposite with down-regulated genes are removed. The final up/down-regulated genes are called invariant differentially expressed genes (iDEGs).

### iDEGs based classification score: gene signature

StepMiner algorithm was used to compute a threshold that separate the high and low values for each gene^55^. To compute the iDEGs signature, the gene expression values will be normalized according to a modified Z-score approach centered around StepMiner threshold (formula = (expr-SThr)/3*stddev). Two iDEGs signature scores: one for Up another for Down is computed by adding together the normalized expression values for every probeset for iDEGs Up/Down. Weighted linear combination (1 for Up, -1 for Down) of these two scores are computed for the final iDEGs signature score. The samples are ordered based on the final iDEGs signature score and ROC-AUC is computed based on the performance of classification of the normal retina vs retinoblastoma samples or two groups of samples with different disease states.

### Visualization and quantification

Gene signature is used to classify sample categories and the performance of the multi-class classification is measured by ROC-AUC (Receiver Operating Characteristics Area Under The Curve) values. A color-coded bar plot is combined with a density plot to visualize the gene signature-based classification. Volcano plot and heatmap were created using python matplotlib package (version 2.1.1). Pathway analysis of gene lists will be carried out via the Reactome database and algorithm^56^.

### Measurement of classification strength or prediction accuracy

Receiver operating characteristic (ROC) curves were computed by simulating a score based on the ordering of samples that illustrates the diagnostic ability of binary classifier system as its discrimination threshold is varied along the sample order. The ROC curves were created by plotting the true positive rate (TPR) against the false positive rate (FPR) at various threshold settings. The area under the curve (often referred to as simply the AUC) is equal to the probability that a classifier will rank a randomly chosen disease samples higher than a randomly chosen healthy samples. In addition to ROC AUC, other classification metrics such as accuracy ((TP + TN)/N; TP: True Positive; TN: True Negative; N: Total Number), precision (TP/(TP+FP); FP: False Positive), recall (TP/(TP+FN); FN: False Negative) and f1 (2 * (precision * recall)/(precision + recall)) scores were computed. Precision score represents how many selected items are relevant and recall score represents how many relevant items are selected. Fisher exact test is used to examine the significance of the association (contingency) between two different classification systems (one of them can be ground truth as a reference).

### Software Code and Reproducibility

Instructions for how to analyze dataset is available at http://hegemon.ucsd.edu/eye/. Bash, perl and python scripts for reproducing the figures and analyses can be downloaded. All datasets used in this paper are available in Gene Expression Omnibus (GEO) website.

### RNA extraction and Real time PCR

Total RNA was extracted from tissues and cells by using TRIzol reagent (Invitrogen) as per manufacturer’s guidelines. The extracted RNA was resuspended in MilliQ water with RNAse inhibitors (Sigma, USA) and the quality was assessed on agarose gel. 1µg of RNA was converted into cDNA using iScript cDNA synthesis kit (Biorad) according to the manufactures protocol. RT-qPCR was done using GoTaq qPCR master mix (promega) with PCR conditions: 50 °C for 2 minutes, 95 °C for 10 minutes, 95 °C for 30 seconds, 60 °C for 1 minute (step3-4 for 40 cycles) and 95 °C for 2 minutes. Data were normalized against the Ct values of the house-keeping gene GAPDH in each sample. The primers used for real time PCR are shown in Supplementary Table 2. The PCR products were detected with CFX96 Touch Real time PCR detection system (Biorad) and analyzed with CFX Maestro software (Biorad).

### 4.5. Western Blotting

Cells collected was dissolved in RIPA (Radioimmunoprecipitation assay) buffer (EMD Millipore Cat No-20-188), sonicated in ice for 15s, centrifuged at 13,000 x *g* for 10 min at 4 °C and the supernatant was collected. Proteins were estimated using BCA protein assay kit (Thermo Fisher Cat No-23227). For Western blot, equal concentration of cell lysates was loaded onto the gel and separated on a sodium dodecyl sulphate polyacrylamide gel (SDS-PAGE) at 100V in electrophoresis buffer (25mM Tris, 190mM Glycine and 0.1% SDS). The proteins were separated and transferred to PVDF membrane (GE Healthcare Cat No-10600023) using semi-dry transblot apparatus (Hoefer) at 1.50 mA/cm^2^. The membrane was then blocked for 1 h at 25 °C in 5% (w/v) non-fat dry milk powder (NFDM) in TBST (20mM Tris-HCl pH 7.5, 150mM NaCl and 0.1% Tween 20). It was washed with TBST, incubated at 4 °C overnight with the β-Catenin (Cell signaling and Technology Cat No-9961) and β-actin (Santa Cruz Cat No-sc47778) antibodies. After overnight incubation, the membrane was washed thrice for 5 min with TBST and further incubated in corresponding HRP-conjugated Anti-rabbit and Anti-mouse secondary antibody). The secondary antibodies used were diluted to 10,000-fold in 5% NFDM (w/v) in TBST. After incubation, the membrane was again washed thrice for 5 min with TBST. HRP activity was detected using HRP substrate (Bio-Rad Cat No-1705061) in Bio-Rad Gel Documentation system (Protein Simple).

### Immunofluorescence

Cells collected were fixed in 4% Paraformaldehyde and washed with Phosphate buffered saline pH 7.2 (PBS). Cells were permeabilized with 0.5% triton-100 followed by PBS wash. The cells were then blocked with 1% BSA prior to overnight incubation with primary Antibody. The detection was done using Cy3.5 secondary antibody and counterstained with Hoechest (Thermo fisher Cat No-33342). Fluorescence microscope (Observer Z1, Carl Zeiss) was used for imaging.

### Cell viability assay

Y79 cells were seeded in 96 well plate to a cell density of 6000/well 24hrs prior to the experiment. The NSC663284 inhibitor (Tocris Biosciences, United Kingdom) was added at increasing concentrations from 50-500nM to the cells in triplicates. The samples were incubated at 37°C with 5% CO2 in a humidified incubator for 24h. The sensitivity of the cells to the inhibitor was evaluated using the colorimetric MTT assay. A multi-well scanner (Spectramax, molecular devices, USA) was used to measure the absorbance at 570 nm wavelength. The untreated/control cells were assigned a value of 100%. Cell survival/viability was calculated by the following equation: (Test OD/Control OD) X 100%.

To determine the live/dead cells, a cocktail of Calcein-AM and ethidium homodimer purchased from Molecular probes was used (4 µM each per sample) and imaged using fluroscence microscope with suitable filters.

### Flow cytometry analysis

To perform the cell cycle analysis Y79 cells were seeded in 6-well plates 24 hrs prior to the treatment of the inhibitor (100nM and 200nM) and incubated again for 24hrs. Post incubation, cells were retrieved and washed twice with cold PBS, fixed with 70% cold ethanol and stored at-20^0^C until used. The cells were thawed on ice and washed twice with PBS and incubated with RNase (10μg/ml) and Propidium Iodide (PI-50μg/ml) at 37°C for 3 hrs. Post incubation, cells were washed twice with PBS, resuspended in sheath fluid and acquired through Flow cytometer (BD, USA). All experiments were performed in triplicates.

### Animal Studies

The animal study has been approved by the Syngene, Institutional Animal Ethics Committee (IAEC Protocol Approval No: SYNGENE/IAEC/537/08-2014) where the study was conducted. Athymic Nude-Foxn1^nu^ female mice with 5-6 weeks of age were injected with 1x 10^7^ of Y79 cells/animal. The cells were allowed to grow until the tumor attained ∼125mm^3^. The mice were stratified into three groups of 8-10 animals: vehicle control, low dose (2.5mg/kg), high dose (5 mg/kg) of NSC663284 CDC25 inhibitor. The mice in the low dose group were injected with the inhibitor every 2 days and in the high dose group, they were injected with the drug after every 4 days. The two regimes, low dose (with frequent injections of drug) and high dose (with less frequency of drug injections) were studied for the drug efficacy studies. Tumor volumes and body weights were calculated after every three days during the study period. The tumor volumes were measured using a Vernier caliper, The length (L) and width (W) of the tumors were measured and tumor volume (TV) was calculated using the following formula: Tumor Volume (mm^3^) = L x W^2^/ 2, Where, L = Length (mm); W = Width (mm). The mice were followed for three weeks and euthanized upon completion of the study period.

### Immunohistochemical and hematoxylin/eosin (H and E) staining

Immunohistochemical and H and E staining were performed on the tumor tissue sections from the mouse xenograft model as previously described [19]. Anti-Ki 67 was purchased from Sigma, USA and used at a dilution of 1:25 for the IHC application.

### Statistical analyses

The statistics for the Cell cycle assay and cell viability assay was performed with Student t-test, and the values with p<0.05, were considered significant. All experiments were performed at least three independent (n=3) times. The statistical significance of the differences was analyzed by paired student “t” test and one-way analysis of variance (ANOVA) followed by Bonferroni Post-hoc test using Graph Pad software. Asterisks *, **, and *** denote a significance with p-Values <0.05, 0.01, and 0.001 respectively.

## Results

### Invariant Differential Expression Analysis (iDEA) identifies invariant differentially expressed genes (iDEGs)

Traditional Differential Expression Analysis (DEA) often suffer from reproducibility problems across independent datasets^11^. The results of this analysis are heavily controlled by the pvalue threshold that is typically adjusted after multiple hypothesis correction^1^. Neither sample size nor pvalue threshold can resolve the reproducibility issues. We have developed a new computational approach called invariant differential expression analysis (iDEA) that has the potential to resolve this reproducibility issues. iDEA performs traditional DEA followed by a Boolean analysis-based filtering step (Fig. 1B). To achieve robustness, iDEA enforce strong Boolean implication relationships (Fig. S1A-G) across and within down- and up-regulated genes in a large diverse gene expression dataset (Fig. 1C, S1I). To perform iDEA on human retinoblastoma we used two independent datasets GSE125903 (2 RT, 7 RB) and GSE97508 (3 RT, 6 RB). First step is to perform a traditional DEA which results in 639 up- and 981 down-regulated genes in GSE125903, and 490 up- and 1139 down-regulated genes in GSE97508 (Fig. 1D, S1H). The DEA analysis on these two cohorts shared 359 down- and 209 up-regulated genes. These genes are filtered using three different types of Boolean implication relationships in a large diverse human microarray dataset GSE119087 (n = 25,955; Affymetrix Human U133 Plus 2.0)^16^. This dataset includes diverse tissues, cancers, diseases and cell lines. We expect that up-regulated genes should have Boolean equivalent relationships with each other. Similarly, down-regulated genes should also have Boolean equivalent relationships with each other. The relationship between an up-regulated gene A and a down-regulated gene B are expected to be A high => B low or A Opposite B Boolean implication relationship. iDEA performs these three filtering steps to identify iDEGs which are supposed to be fundamental gene regulatory changes that hold across almost all tissues regardless of their disease state. Therefore, reproducibility across dataset are expected to improve with iDEA. Finally, 14 genes down-(Supplementary Table 5) and 70 genes up-regulated (Supplementary Table 6) that are shared between GSE125903 and GSE97508 (Fig. 1E). Top five genes downregulated are GABRA1, OPCML, B3GAT1, RIBC1, and FRMPD4 (Fig. 1F). Top five genes upregulated are MYBL2, FAM72A, CKAP2L, UBE2C, and DEPDC1B (Fig. 1F).

### Invariant differentially expressed genes (iDEGs) robustly classify retina disease states and reveals mechanism of cancer resistance

To classify retina disease states, both up- and down-regulated genes in the iDEGs are used to compute a gene signature based on weighted linear combination of the normalized gene expression values (See methods). Gene signature is used as a score to classify the sample categories. The performance of this classification is measure using Receiver operating characteristic (ROC) area under the curve (AUC). ROC-AUC close to 1 is considered good classification and 0.5 is considered the worst possible classification. Performance of iDEGs (Boolean), iDEGs/up, iDEGs/down is compared against other gene signatures from the literatures in two human datasets (GSE87042, GSE24673)^39,57^ and two mouse datasets (GSE29686, GSE86372)^58,59^ by computing the average of the ROC-AUC values (Fig 2A). Up regulated iDEGs (Boolean/Up) performed the best followed by combined iDEGs signature. Classification of retina vs RB using the up-regulated iDEGs is validated in four independent retina and retinoblastoma datasets (GSE97508, GSE125903, GSE87042, GSE110811; ROC-AUC > 0.85; Fig 2B)^57,60,61^. If a gene signature classifies retina vs RB robustly, it may not classify severity of RB. However, we observed that up regulated iDEGs (Boolean/Up) classify severity of RB in three independent datasets (GSE59983, GSE110811, GSE29686; ROC-AUC > 0.83; Fig 2C)^59,60,62^. Up regulated iDEGs also predicted therapeutic effects of chemotherapy, developmental stages of retina, and various Rb mutant retina (GSE24673, GSE74181, GSE86372; ROC-AUC = 1.0; Fig 2D)^39,58,63^. We studied three different RB cell lines (resistant NCC; sensitive Weri and Y79) that model resistance to cell cycle inhibitors. Up regulated iDEGs based gene signature placed NCC close to normal and both Weri and Y79 far from normal retina. We performed a high-resolution differential expression analysis to identify genes that may distinguish NCC from Weri and Y79 (Fig 2E). 436 genes were differentially expression in both Retina vs NCC and Weri/Y79 vs NCC after removing the retina vs Weri/Y79 background DEGs (Fig 2F). Reactome pathway analysis of these 436 genes revealed Wnt/β-catenin pathway significant (Fig 2G).

**Figure 2:**
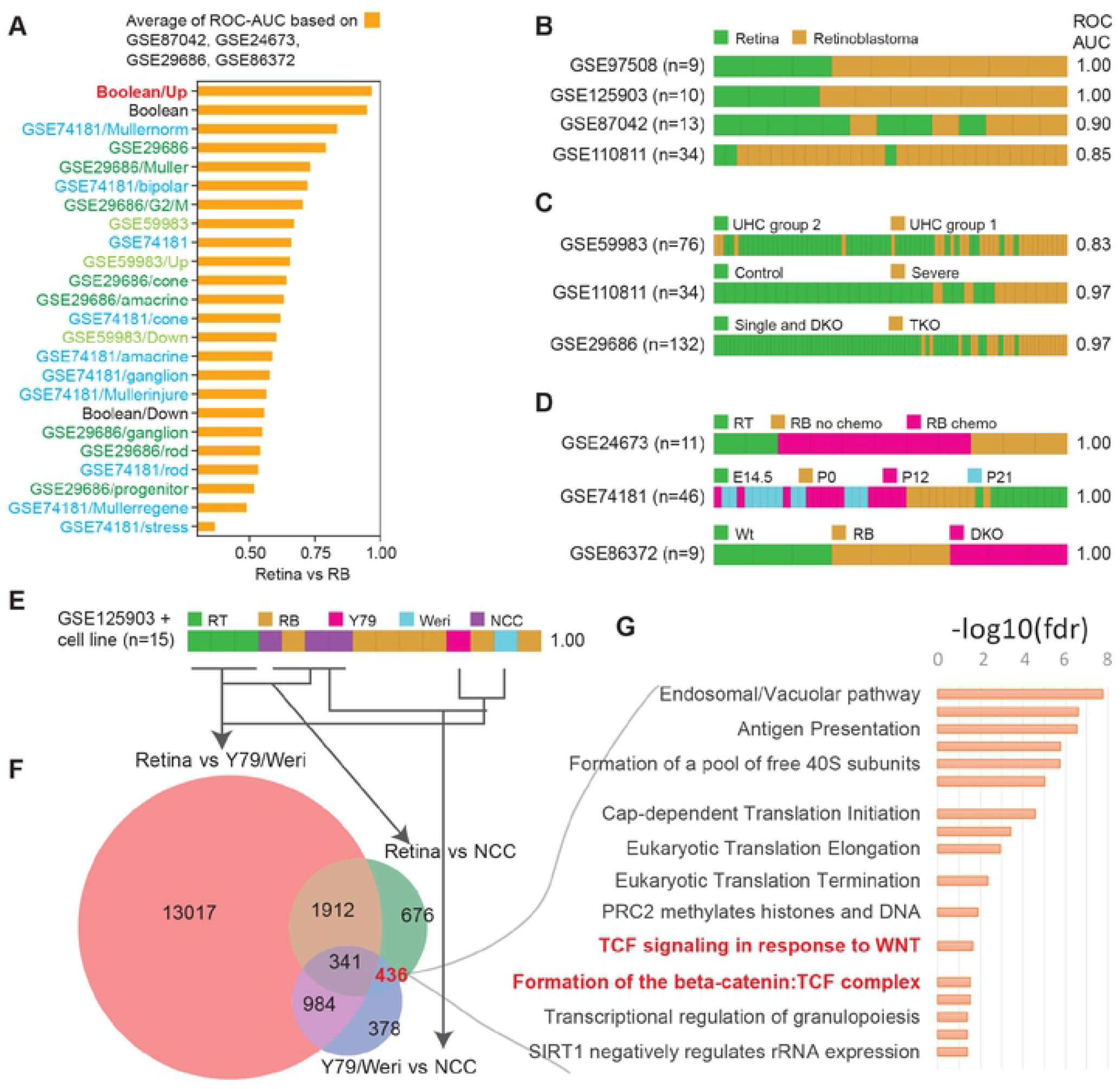
RB gene signature reveals mechanism of cancer resistance. (A) Boolean invariant differentially expressed genes between normal retina and retinoblastoma: Boolean/Up, Boolean/Down and Boolean (Combined Up and Down) are compared against other gene lists from literature. Average ROC-AUC based on two human datasets (GSE87042, GSE24673) and two mouse datasets (GSE29686, GSE86372) is used to identify the top scoring gene signature. (B) Validation of Boolean/Up gene signature in four independent retina and retinoblastoma datasets. (C) Boolean/Up gene signature discreminate tumor phenotypes in two human (GSE59983; UHC group 1 vs 2, GSE110811; differentiation status: Severe vs non-severe) and one mouse dataset (GSE29686; single and double knockout DKO vs tripple knockout TKO). (D) Boolean/Up gene signature discreminate chemo-treated samples (GSE24673), mouse retina at different developemental stages (GSE74181), mouse retina genotypes (GSE86372): wild type (WT) vs single RB knockout vs double knockout (DKO). (E) Boolean/Up gene signature placed NCC cell lines close to retina and Y79/Weri cell lines far from retina. (F) 436 genes are differentially expressed between retina and NCC after removing background retina vs cell line differences. (G) Pathway analysis of 436 genes identified enrichment of Wnt/TCF/beta-catenin signaling.

### Transcripts levels of β-catenin and CDC25 are strongly correlated in both human and mouse datasets

We asked whether β-catenin (CTNNB1) transcripts levels are strongly associated with cell cycle. We checked the relationship between CTNNB1 and CDC25A/B/C in both human and mouse datasets. The mouse dataset included RNASeq from normal mouse retina (GSE87043, n=8). To get a diverse human dataset, we performed xenografts and cultured in two different conditions (See supplementary methods) using NCC cell lines. Two different culture conditions were used to grow NCC cell lines: standard tissue culture plates and graphene sponge. Graphene sponge were synthesized from nickel foam using standard techniques (See supplementary methods; Figure S4). NCC cell line was cultured in a 3D environment in graphene sponge. Finally, the human dataset included normal retina, retinoblastoma, xenografts, and RB cell lines (Total n = 22; GSE125903, retina n=2, RB, n=7; NCC Xenograft n=3; NCC tissue culture n=3; NCC graphene sponge n=3; GSE141327, Weri n = 2, Y79 n = 2). In both human and mouse dataset we observed a strong correlation between CTNNB1 and CDC25A/B/C (Fig 3A-B). We also observed that both Weri and Y79 has higher expression patterns for CDC25A/B/C compared to NCC and normal retina. Therefore, we predicted that Weri and Y79 may have higher levels of cell cycle activity compared to NCC. We validated the elevated expression of CDC25 transcripts in additional RB cohort using qPCR. We observed that CDC25A (p<.05), CDC25B (p<.005), CDC25C(p<.05) were significantly up regulated in human RB samples (Fig S3A).

**Figure 3:**
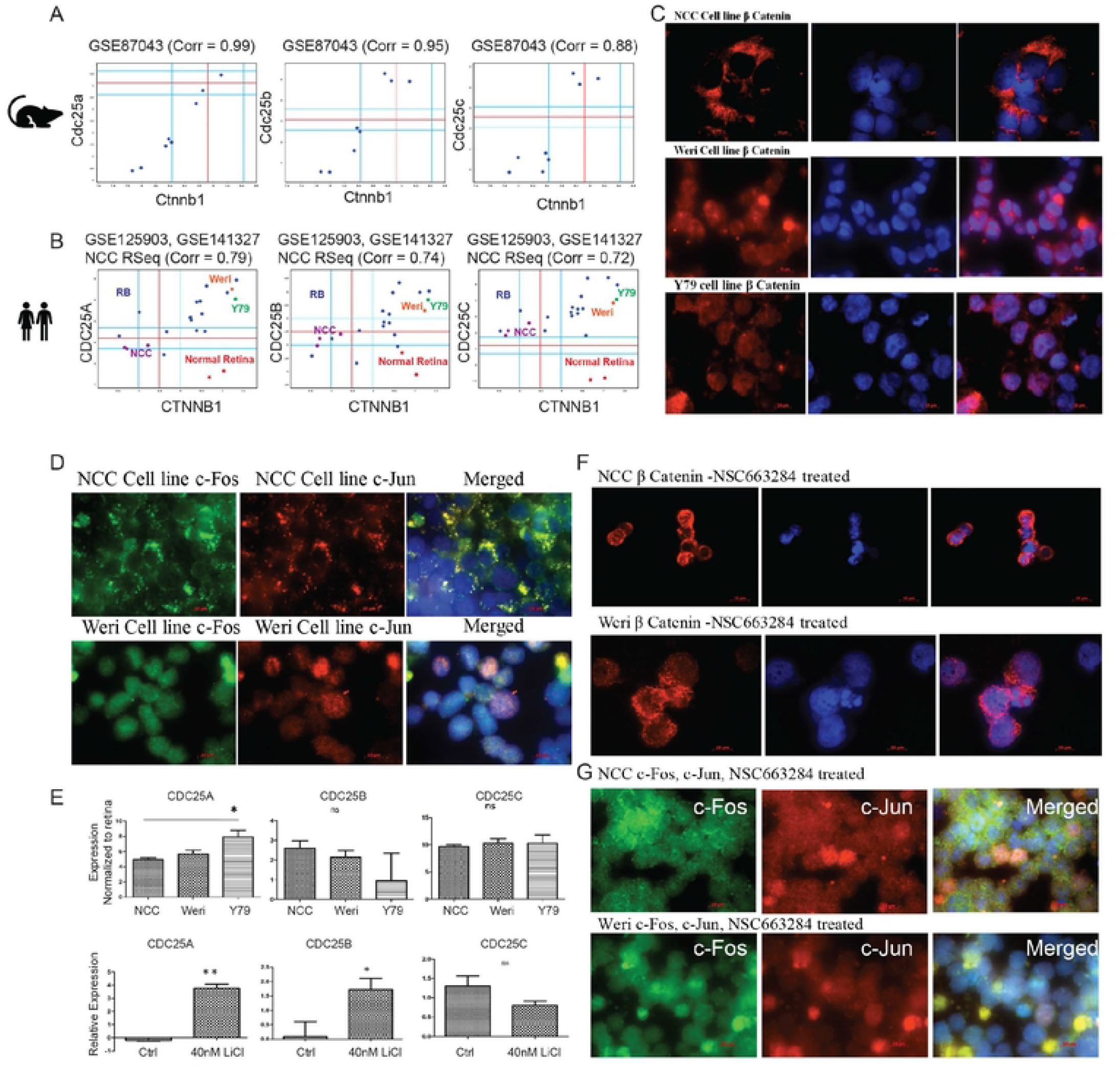
Beta-catenin pathway mediate cancer resistance. (A) Correlation analyses between CDC25 and beta-catenin mRNA expression in normal mouse retina (GSE87043, n=8). Pearson’s correlation coefficient is displayed at the top of each scatterplots. (B) Correlation analyses between CDC25 and beta-catenin in human retinoblastoma, xenografts, and RB cell lines (GSE156657: GSE125903, RB, n=7; NCC n=3; GSE141327, Weri n = 1, Y79 n = 1). Normal human retina (GSE125903, n=2) is displayed in the scatterplots. (C) beta-catenin localization studies using Immunofluroscence in RB cell lines: NCC (top), Weri (middle) and Y79 (bottom). (D) c-Fos and c-Jun localization using immunofluorescence in NCC and Weri cell lines. (E) Localization of beta-catenin in NCC (top) and Weri (middle) after CDC25 inhibitor (NSC663284) treatment. (F) Localization of c-Fos (left) and c-Jun(middle) after CDC25 inhibitor (NSC663284) treatment. (G) Relative expression of CDC25 transcripts in RB cell line after normalized to retina in NCC, Weri, and Y79 (top panels). Effect of LiCl treatment on CDC25 transcripts in NCC cell line (bottom panel).

### Resistant NCC Rbc51 cell line exhibits Differential Wnt signalling and interacting partners compared to sensitive Y79 and Weri cell lines

CDC25s are phosphatases which control cyclin-dependent kinases and play prominent role in cycle progression. β-catenin is known as the central molecule in Wnt pathway signalling. Therefore, we investigated if these two prominent molecules that effected cell cycle, influenced mutual gene expression in RB tumours. β-catenin exists in three different subcellular locations as membrane bound, cytosolic, and nuclear locations. Further, experiments were undertaken in cell culture models to understand the expression and localization of β catenin. RB cell line such as NCC, Weri and Y79 cells were analysed for its expression and localisation. Weri and Y79 showed complete nuclear localization of β-catenin whereas in NCC Rb51 has a membrane localization (Fig 3C). In different cancer lines, β-catenin has been showed to have physical association with c-Fos and c-Jun^64^. Hence, we further analyzed the localization of these proteins in NCC Rb51 and Weri through co immunostaining. Interestingly, both c-Fos and c-Jun showed differential localization in both the cell lines (Fig 3D). From these data we can observe that the localization of the transcription factors such as β-catenin, c-Fos and c-Jun are different in NCC Rb51 compared to Weri/ Y79 cell lines.

CDC25 gene expressions were higher in Weri and Y79 compared to NCC Rb 51 (Fig 3B). Wnt/Beta catenin pathway is known to be active in most cancers and β-catenin localisation is observed in nucleus in Weri and Y79 cell line (Fig 3C). However, membrane localization observed in NCC Rb 51cell line was intriguing. Therefore, we treated NCC cell line with 40mM of Licl which is an agonist of the canonical wnt signaling and is known to induce nuclear localization of β catenin, which could alter CDC25 expression. The CDC25 A (p<.05), and CDC25B (p<.005) gene expressions were significantly upregulated whereas CDC25 C changes were not significant. (Fig 3E). Our data suggest that the localization of β catenin regulates the CDC25 transcripts expression in this human Rb cell line.

### β-catenin localization governs efficacy of CDC inhibitors

Next, we investigated if CDC25 inhibitor alters the localization of β catenin in human Rb cell line. In NCC Rb 51 cell line treatment with CDC25 inhibitor NSC663284 changed the β catenin localization from plasma membrane to cytosol (Fig 3F). Additionally, the inhibitor treatement induced localization changes of its interacting partners such as c-Fos and c-Jun (Fig 3G) Interestingly, absolute protein levels were unaltered by the NSC663284 treatment (Fig 4A). However, in Weri and Y79 cell line where nuclear localization of β catenin was observed, the inhibitor induced puncta formation indicating cellular apoptosis. Hence the effect of CDC inhibitor for inducing cell death was dependent on β catenin localization. The drug efficacy could be higher in the cells where β catenin is in nucleus. Therefore, the active wnt signaling is needed for the drug efficacy studies.

**Figure 4:**
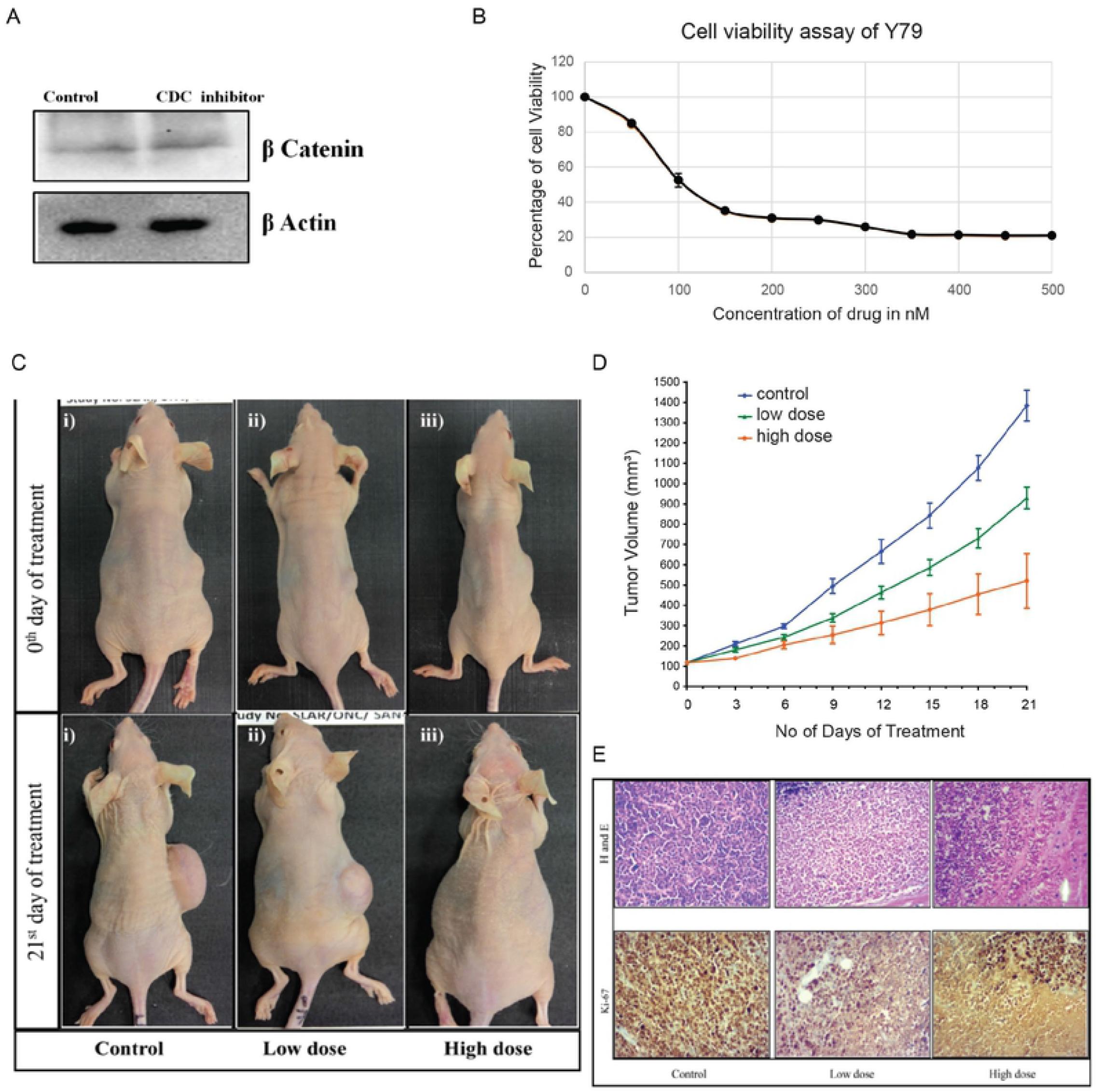
CDC inhibitor blocks growth in beta-catenin active tumor. (A) Immunoblotting of beta-catenin in NCC with and without CDC inhibitor. No change in beta-catenin protein level after CDC inhibitor treatment. (B) MTT assay showing the different concentration of CDC inhibitor and cell viability. (C) CDC25 inhibitor is effective in reducing tumour volume in RB mouse xenograft model. (D) Graphical representation of tumor growth inhibition in mm^3^ by CDC25 inhibitor (NSC663284) in RB mouse xenograft models. The results shown in are the mean ± S.D of the tumor volume for 10 animals in the control group, 8 animals in the low dose group, 7 animals in the high dose group. (E) Immunohistochemical and H&E staining was performed on the tumor tissue sections from the mouse xenograft model.

### CDC25s are effective therapeutic target in β-catenin active RB

Further to validate the effect of CDC25s inhibitor on Y79 cell line (beta-catenin active cell line) we evaluated its effect in mouse xenograft model. To evaluate the effect of the CDC25 inhibitor NSC663284 on the Y79 cell line, different concentration of the CDC25 inhibitor (50 to 500 nM) treatment were used for 24h (Fig 4B). We observed that 100-150nm of NSC663284 CDC25 inhibitor treatment showed nearly 50% viability in Y79 cell lines. The percentage of cell viability decreased with increased inhibitor concentration. We performed the cell cycle analysis to identify the effect of the inhibitor on the Y79 cells. There was significant (p≤0.05) decrease in the G0-G1 and significant increase (p≤0.05) in the G2-M and S phase arrest of the cells when compared to control, at 100nM concentration of the inhibitor. In the 200nM inhibitor treated cells there is a significant decrease (p≤0.005) in the G0-G1 phase and S (p≤0.05) phase whereas significant increase (p≤0.001) in the G2-M phase of the cells was observed when compared to the untreated cells. Increasing the inhibitor concentration from 100nM to 200nM resulted in a significant increase (p≤0.05) in the G2-M phase and significant decrease (p≤0.05) in the S phase (Fig S3).

Mouse xenograft models of RB was generated injecting Y79 cells into the nude mice subcutaneously to develop measurable tumors of 100 mm^3^. The mice were treated with low dose (2.5mg/kg injected every other day for 3 weeks) and high dose (5mg/kg injected every 4 days for 3 weeks) of the inhibitor. We observed that a significant reduction in the tumor volumes with both dosages conditions but significantly more tumor reduction (62%, p≤0.05) was observed in mice treated with the high dosage compared to mice treated with low dosage (Figure 4C and 4D). There was no significant (p>0.2) decrease in the body weight during the treatment regime. We elucidated the effect of the inhibitor on the other organs such as the liver, kidneys, lungs and spleen of the mice and observed no profound adverse effect.

Immunohistochemistry (IHC) was performed on the xenograft tumors using anti Ki67 antibody which is a proliferative marker to assess the effect of the inhibitor on the tumors. We observed that mice that were not treated with the inhibitor exhibited higher expression of the Ki67 protein in the tumor tissues. In the low dose inhibitor group decreased expression of Ki67 protein was observed. We observed an increase in necrotic cells. Furthermore, in the high dose inhibitor group higher number of necrotic cells was observed in addition to the lower expression levels of Ki67 protein (Figure 4E).

### iDEGs are robustly associated with outcome in breast and prostate cancer

iDEGs helped elucidate the mechanism of drug sensitivity in RB cell lines. Since iDEGs are supposed to be invariant, they represent common pathway that are fundamental cell cycle activity in diverse tumors. To test if iDEGs can be translated to other tumor types, we investigated how strongly iDEGs are associated with outcome. Cell cycle activity is not universally prognostic in all tumors. Cell cycle activities are known to be prognostic in breast cancer and prostate cancer. However, prognostic significance of cell cycle is not established in colorectal cancer, pancreatic cancer, and lung cancer. We predicted that iDEGs signature may be prognostic in breast cancer and prostate cancer, but it may not be prognostic in colorectal cancer, pancreatic cancer and lung cancer. We tested iDEGs signature in two independent breast cancer cohorts: pooled cohort of GSE2034+GSE2603+GSE12276 (n=572) and METABRIC (n = 2136). iDEGs signature is computed using linear combination of normalized Z-scores around StepMiner threshold (See supplementary methods). Final iDEGs signature score is divided into high and low values using StepMiner. In both breast cancer cohorts high iDEGs signature is associated with worse outcome (GSE2034+GSE2603+GSE12276, p = 0.016 in ESR1 low, p < 0.001 in ESR1 high; METABRIC, p = 0.013 in ESR1 low, p = 0.0026 in ESR1 high; Fig 5A-B). Univariate and Multivariate analysis (Fig 5B) in METABRIC cohorts suggest that iDEGs is better than differentially expressed genes (DEGs) in the individual RB datasets (GSE97508 and GSE125903). High iDEGs signature score was also significantly associated with worse outcome in two independent prostate cancer cohorts (GSE16560, p < 0.001; GSE21034, p < 0.001; Fig 5C). However, association with outcome was not significant in pancreatic cancer, colorectal cancer and lung cancer (Fig S5). Only one lung cancer cohort (GSE68465; n=462) shows significant association with outcome (Fig S5C). These data suggest that iDEA significantly improves differential expression analysis.

**Figure 5:**
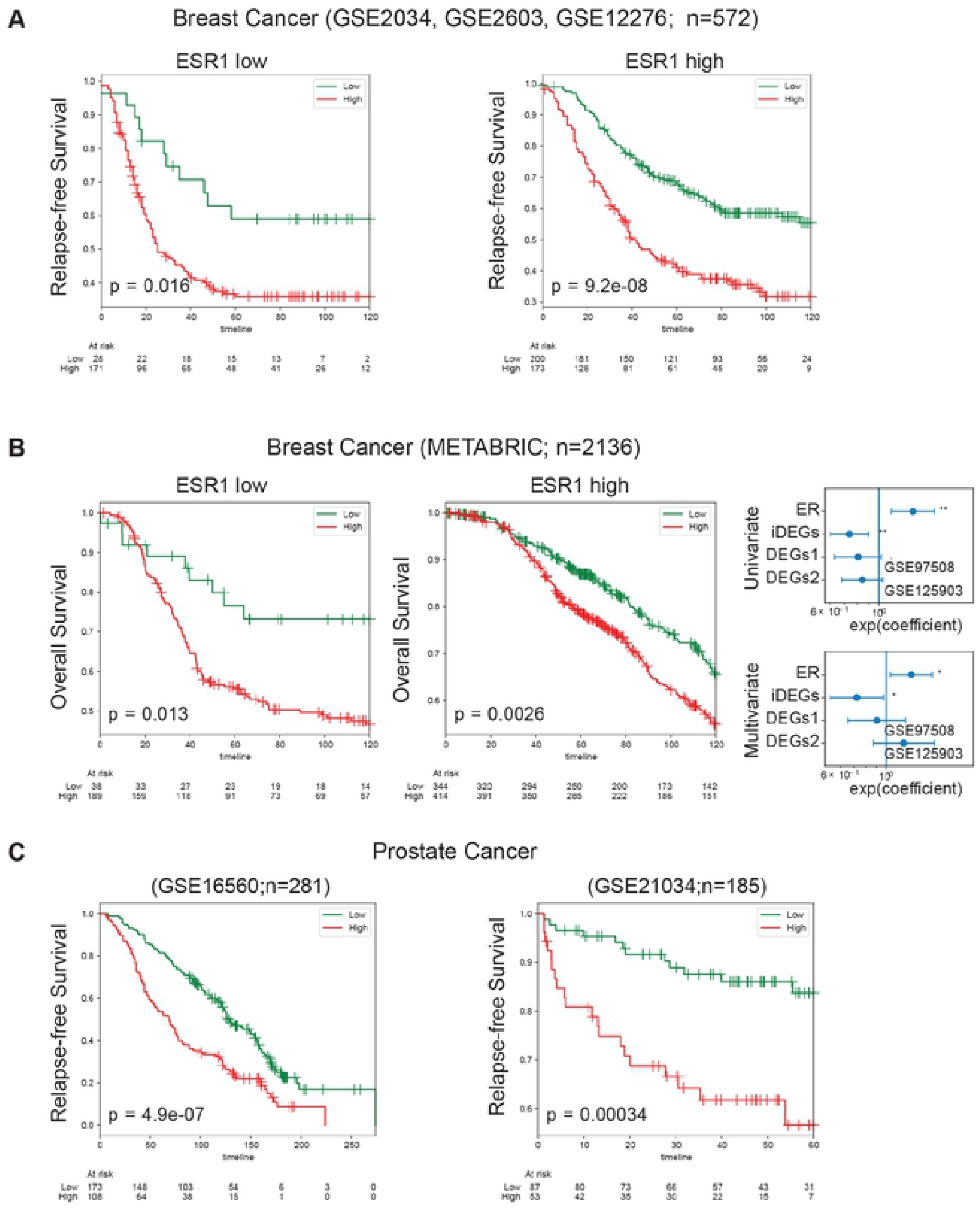
iDEG signature is prognostic in breast and prostate cancer. (A) Three breast cancer datasets (GSE2034, GSE2603, and GSE12276) are combined to create this cohort with consistent relapse-free survival annotations. iDEG signature is used to classify breast cancer patient samples into high and low subgroups in both ESR1 low (left) and ESR1 high (right) tumors. (B) METABRIC breast cancer samples annotated with overall survival is subjected to the same analysis in panel A. (C) iDEGs signature is used to classify prostate cancer samples into high and low subgroups in two independent datasets (GSE16560, left; GSE21034, right). (A-C) Kaplan-Meier analysis in performed using python lifelines package and p values are computed using log-rank test. Both analyses are independently verified using R statistical software. Univariate and Multivariate Cox-proportional hazard analysis is performed in METABRIC dataset (Panel B) to compare between iDEGs and DEGs.

## Discussion

It is well known that differential expression analysis (DEA) suffer from reproducibility issues because of diversity in patient cohorts. Here we introduce a new method called invariant differential expression analysis (iDEA) that use strong Boolean implication relationships in independent largescale pooled cohorts of gene expression datasets to improve reproducibility. Despite smaller cohorts of available retinoblastoma datasets, iDEA was able to identify an invariant differentially expressed genes (iDEGs) that is successfully validated in multiple independent cohorts of RB datasets. It is also surprising to see that the results are still relevant in diverse cancer datasets including breast and prostate cancer. iDEA enabled discovery of Wnt signalling as a major pathway that mediate sensitivity to cell cycle inhibitors. The gene annotations for the iDEGs revealed that many of the genes involved in the DNA synthesis, DNA repair, nucleosome assembly and cell cycle. The reactome pathway analysis identified G1-S phase of the cell cycle as a major deregulated pathway in RB tumour progression. Since, iDEGs can classify RB cell lines that are known to have sensitivity to cell cycle drugs, we focused our analysis on this process for identifying potential mechanism of action for the drugs sensitivity. Among the cell cycle genes that classify normal/tumors in both mice and human, CDC25 molecules emerged as therapeutic targets. CDC25 has been reported to be overexpressed in many human cancers^65^. However, its association with clinical prognosis has been difficult to assess presently^66^. The mechanism by which it becomes dysregulated is still unclear since its regulation takes place at multiple stages involving promotor methylation, transcript and protein regulation. Additionally, post translational regulation was also observed. CDC25 transcript level has been reported to be upregulated in SV40 transformed and adenovirus infected fibroblast cell^67^, suggesting CDC25 promoters are specifically targeted by activated transcription factors. Various transcription factors have been known to regulate CDC25 expression such as E2F1/2/3,Stat3,Foxm1 and c-Myc^66^.

Wnt signaling pathway which has been primarily involved in development and evolution process has been recently linked with cell cycle regulation. As reported earlier most of the wnt beta catenin component pathways has been found in centrosomes^68^. These may facilitate the pathway of proximity of regulators. However in most of the cancers Wnt pathway has been reported to be activated showing nuclear localization of β catenin^69^. We observed that the Y79 and Weri cell lines had nuclear localization of β catenin indicating that wnt signalling may be active in these cell lines. However, membrane localisation in aggressive NCC RB51 cell line indicates that wnt signalling is not active in NCC Rb cell line and after using an inhibitor for canochial wnt signalling Beta catenin localised to nucleus activating wnt signalling. Since, the growth pattern and drug sensitivity are different between these two cell lines, we investigated the relationship between CDC25A/B/C and β catenin. The bioinformatic analysis indicated a positive correlation between CDC25A/B/C expression and β catenin in both mouse retina and human RB samples.

CDC25 belongs to the family of proteins with conserved dual specific phosphatase activity. It activates CDKs, which in turn regulates the cell cycle progression^70^. We observed increased CDC25A and CDC25C gene expression in our transcriptomics data, whereas higher expression of CDC25A and CDC25B at the protein level was reported earlier in RB tumors. Therefore, it is reasonable to hypothesize that CDC25 inhibitors may provide therapeutic target in RB. Various CDC25 small molecules inhibitors has been designed with limited success^71^. The sub-group of patients who respond to CDC25 therapy is unknown probably limiting its use as therapy molecules for various cancers. In this study we identified the differential localization of β catenin and its interacting partner c-Fos and c-Jun could play a prominent role in predicting the efficiency of CDC 25 inhibitors. Active wnt signaling in the nucleus of RB cell lines could be used for stratification of patients who may benefit from CDC25B inhibitors. Furthermore, our results may indicate the existence of a feedback loop between β catenin and CDC25, which could play a prominent role in regulating cell cycle progression leading to therapy in RB. Interestingly, NSC663284 (a potent CDC25 inhibitor) mediated Beta catenin localization in RB cells which in turn may regulate CDC25 proteins that controls apoptosis. Further, reduction in G2/M phase and increase in cell cycle arrest was observed in Y79 cells after the inhibitor treatment. This result is consistent with earlier studies that reported elongation of S phase and cell cycle arrest in mouse mammary carcinoma and human pancreatic ductal adenocarcinoma cancer cells after the drug treatment^72^. The reduction of the tumor growth is highly significant in the high dose group compared to low dose treated group. The observed reduction in tumour growth could be due to the decreased proliferation. Immunohistochemical staining of the xenograft tumors using the Ki-67 proliferative marker showed lower expression in the treatment group, which suggests that there is reduction in tumor cell growth. Therefore, a small molecule inhibitor regulating CDC25s expression can effectively induce tumor reduction in mouse xenograft model of RB indicating their prominent role in RB tumor progression. However, wnt signalling is necessary for the efficacy of this inhibitor. The relationship between wnt activation and efficacy of the CDC inhibitors needs further investigation in additional cohorts. In conclusion, CDC25 phosphatase activity inhibition in wnt activated tumours may provide a new therapeutic option for treating RB tumours in pre-clinical animal models. These targeted CDC25 drug molecules could reduce the off-target effects associated with conventional drugs. These could be used in combination with conventional drugs to bring down the dose needed for the therapy.

## Conclusion

We demonstrated that invariant differential expression analysis (iDEA) improve reproducibility of traditional differential expression analysis. iDEA was applied to retinoblastoma datasets to identify an invariant differentially expressed genes (iDEGs) that is successfully validated in multiple independent cohorts of RB datasets. Wnt signalling was found to be a major pathway that mediate sensitivity to cell cycle inhibitors and xenograft retinoblastoma tumors respond to cell cycle inhibitor NSC663284. Since iDEGs are also found to be relevant in breast cancer and prostate cancer, the mechanism of action for cancer resistance may also be relevant in other cancer types. Further studies on this may reveal that wnt/beta-catenin may be a fundamental pathway for cancer resistance.

## Abbreviations

Idea: invariant differential expression analysis
iDEGs: invariant differentially expressed genes
RB: retinoblastoma tumors
Rb: Retinoblastoma protein
RT: retina
ROC: Receiver operating characteristic
AUC: Area under the curve
GEO: Gene Expression Omnibus
BIRs: Boolean implication relationships
BIN: Boolean implication network
NCBI: National Center for Biotechnology Information
EMBL-EBI: European Molecular Biology Laboratory European Bioinformatics Institutes
RMA: Robust Multichip Average
TPM: Transcripts Per Millions

## Ethical Approval and Consent to participate

The present study was conducted at the Medical Research Foundation and Vision Research foundation Sankara Nethralaya, India. The study was approved by the Institutional Ethics Board. Ethics No. 247-2011-P. Normal adult retinas (n=3, age 30 yrs) were obtained from the enucleated eyeballs of cadaveric donors donated to C.U SHAH eye bank, Sankara Nethralaya. A part of the tumor from the enucleated eyeball was used for pathological evaluation and the remaining part was used for gene expression analysis by Q-PCR.

## Consent for publication

Not applicable

## Availability of supporting data

The datasets generated and analysed during the current study are available in Gene Expression Omnibus (GEO) repository under accession number GSE156657. Instructions on how to analyze dataset is available at http://hegemon.ucsd.edu/eye/. Bash, perl and python scripts for reproducing the figures and analyses can be downloaded. All datasets used in this paper are available in Gene Expression Omnibus (GEO) website.

## Authors’ contributions

DS, SE conceived the idea. AC, RD, RR, RS, MS performed experiments. VK, PR provided the RB tumours after informed consent of patient relatives. DS, SP performed bioinformatic analysis. AC, RD, SE and DS wrote the manuscript and all authors edited the manuscript.

## Acknowledgements

We thank Dr Krishnakumar Subramanyan for grading the tumours. We thank Dr J. Narayanan for performing SEM experiments purched from DBT grant (BT/PR/26926/NNT/28?1500/2017). We thank Dr Anup Chugani from Medenome with RNA sequencing experiments.

## Figure legends

**Figure S1:**
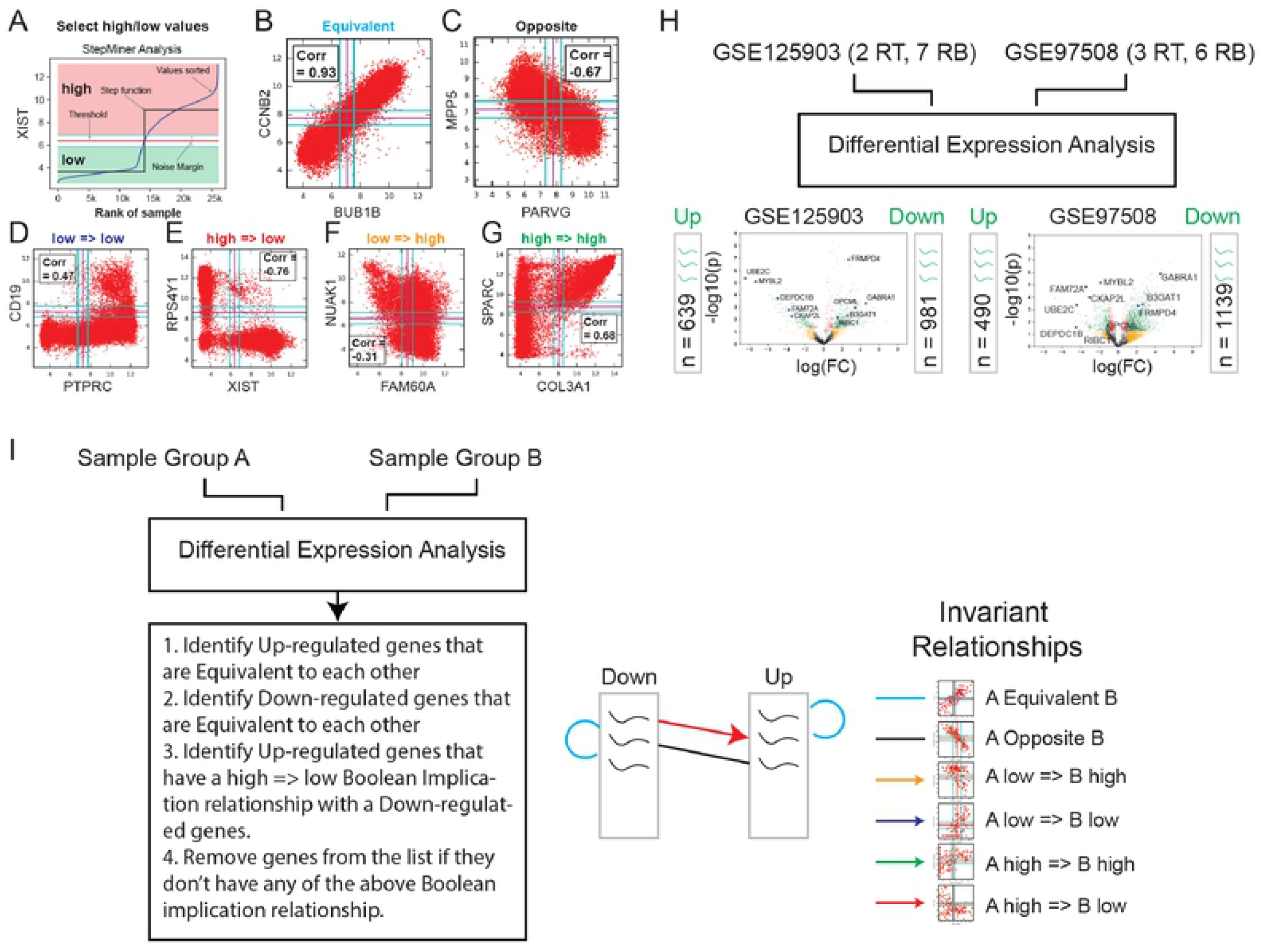
Identification of invariant differentially expressed genes between normal retina and retinoblastoma. (A) Schematic approach to identify high and low expression values (Boolean values) in a global diverse tissue dataset GSE119087 (n = 25,955; Affymetrix Human U133 Plus 2.0). (B-G) Boolean Implication Analysis. (B) Equivalent. (C) Opposite. (D) low => low (E) high => low (F) low => high. (G) high => high. (H) Differentially expressed genes between normal retina and retinoblastoma datasets: GSE125903 (2 control retina vs 7 retinoblastoma; RNASeq) and GSE97508 (3 control retina vs 6 retinoblastoma; microarray). Volcano plots of both datasets. (I) Schematic approach for filtering traditional differential analysis by using Boolean implication relationships (color coded). Pseudocode of the filtering steps are given in the schematic.

**Figure S2:**
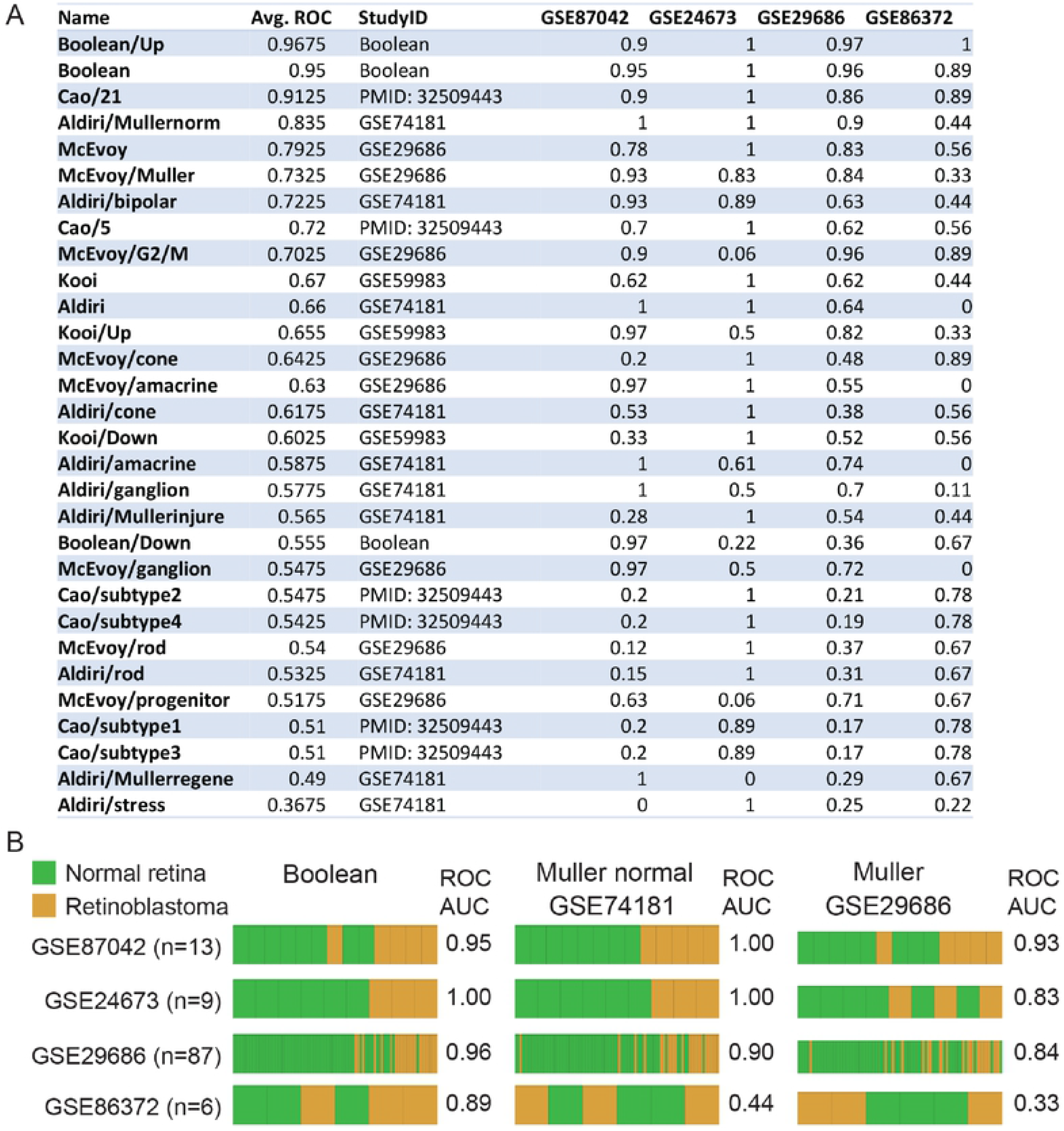
Classification of normal retina and retinoblastoma. (A) Classification of normal retina and retinoblastoma using gene signatures from the literature. ROC-AUC is computed in two human datasets (GSE87042, GSE24673) and two mouse datasets (GSE29686, GSE86372). Average of all four ROC-AUC is computed which is shown in third columns. (B) Barplots to show classification of retina and retinoblastoma in four independent datasets for three selected gene signatures: Boolean, GSE74181/Muller and GSE29686/Muller.

**Figure S3:**
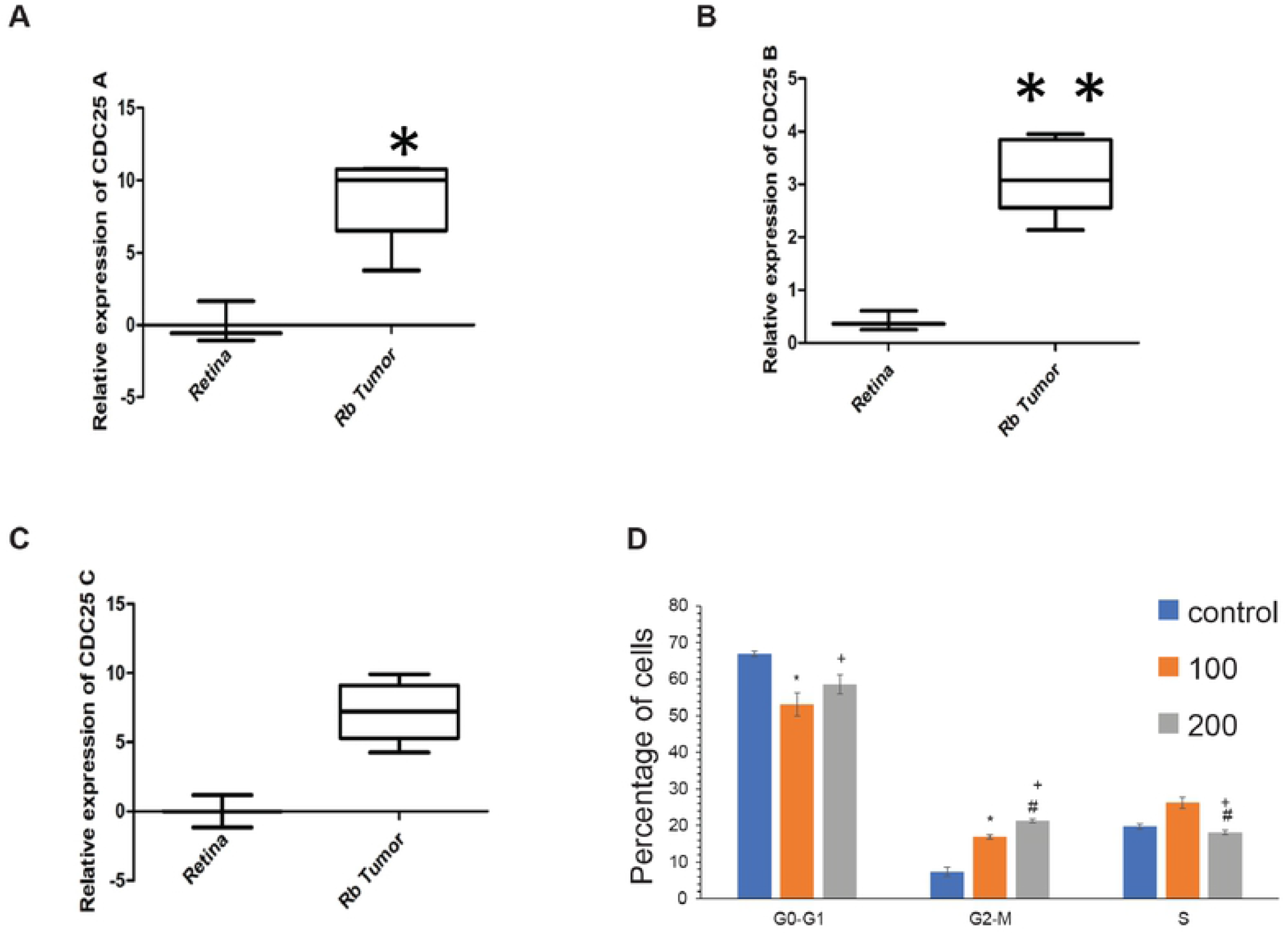
Expression patterns of CDC25 in RB and cell cycle phases. (A-C) qPCR ofCDC2SA/B/C in additional cohorts of 3 retina and 10 retinoblastoma samples. (D) Cell cycle analysis on Y79 cell treated with 100 and 200nM concentration of the inhibitor (NSC 663284). * indicates the significance between Control vs 100nM treatment, + indicates the significance betweenControl vs 200nM treatment, # indicates the significance between 100nM vs 200nM treatments. p<0.05 considered significant.

**Figure S4:**
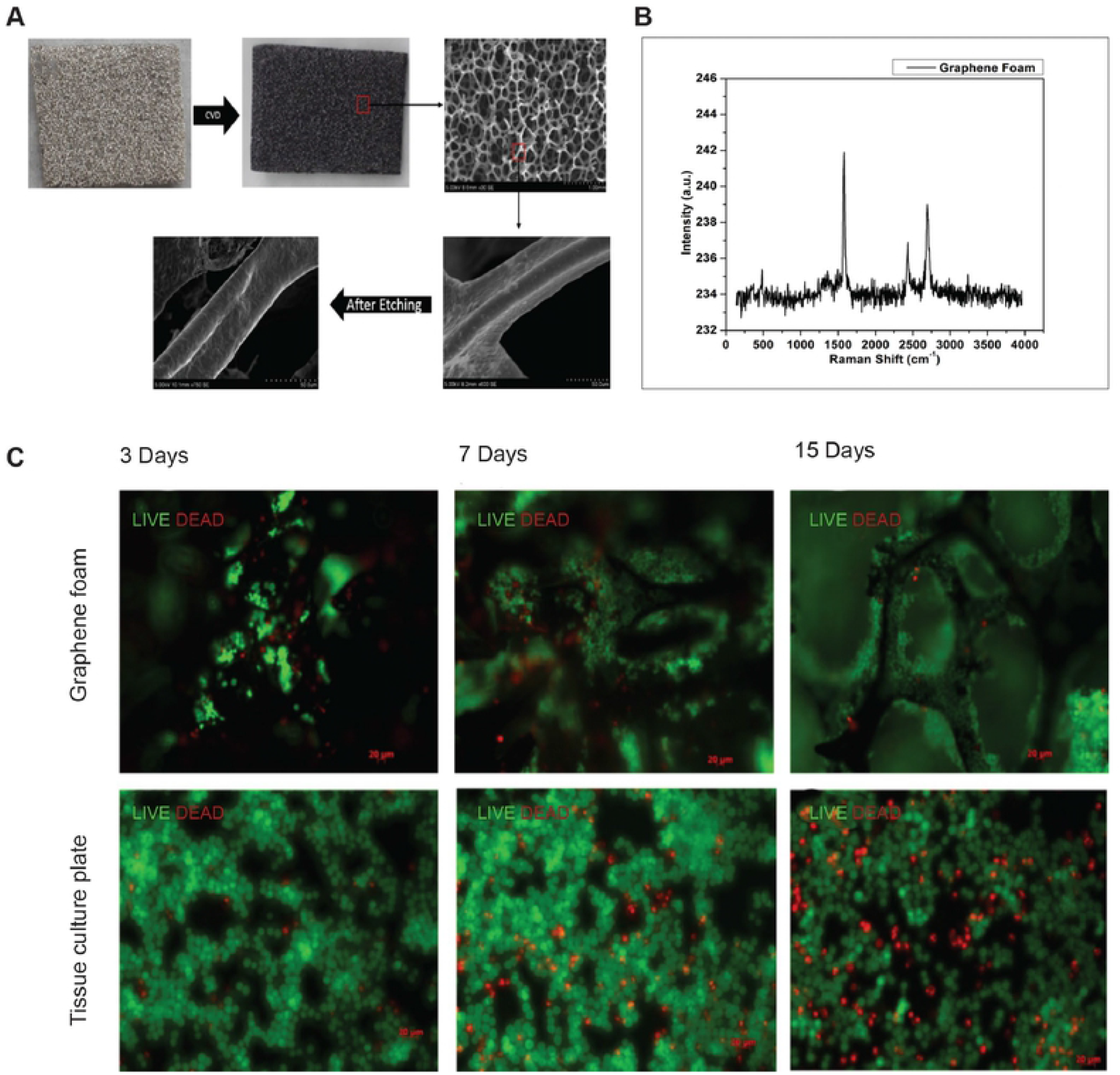
Graphene foam scaffolds synthesis and culture platform. (A) Synthesis of Graphene foam scaffolds using carbon deposition at high temperature on a nickle (Ni) foam followed by Ni etching solution (See supplementary methods). (B) Raman spectroscopy is performed to determine the number and orientation of layers andthe quality of scaffolds. The Raman spectrum indicates graphitic carbon structure. (C) Retinoblastoma cell line NCC-RbC-51 was cultured over graphene sponge and tissue culture plate to understand the heterogeneity of cell growth. Pictures were taken in three different days: 3, 7, and 15. Live and dead cells were visualized using Calcein and ethidium homodimer staining followed by fluorescence microscopy.

**Figure S5:**
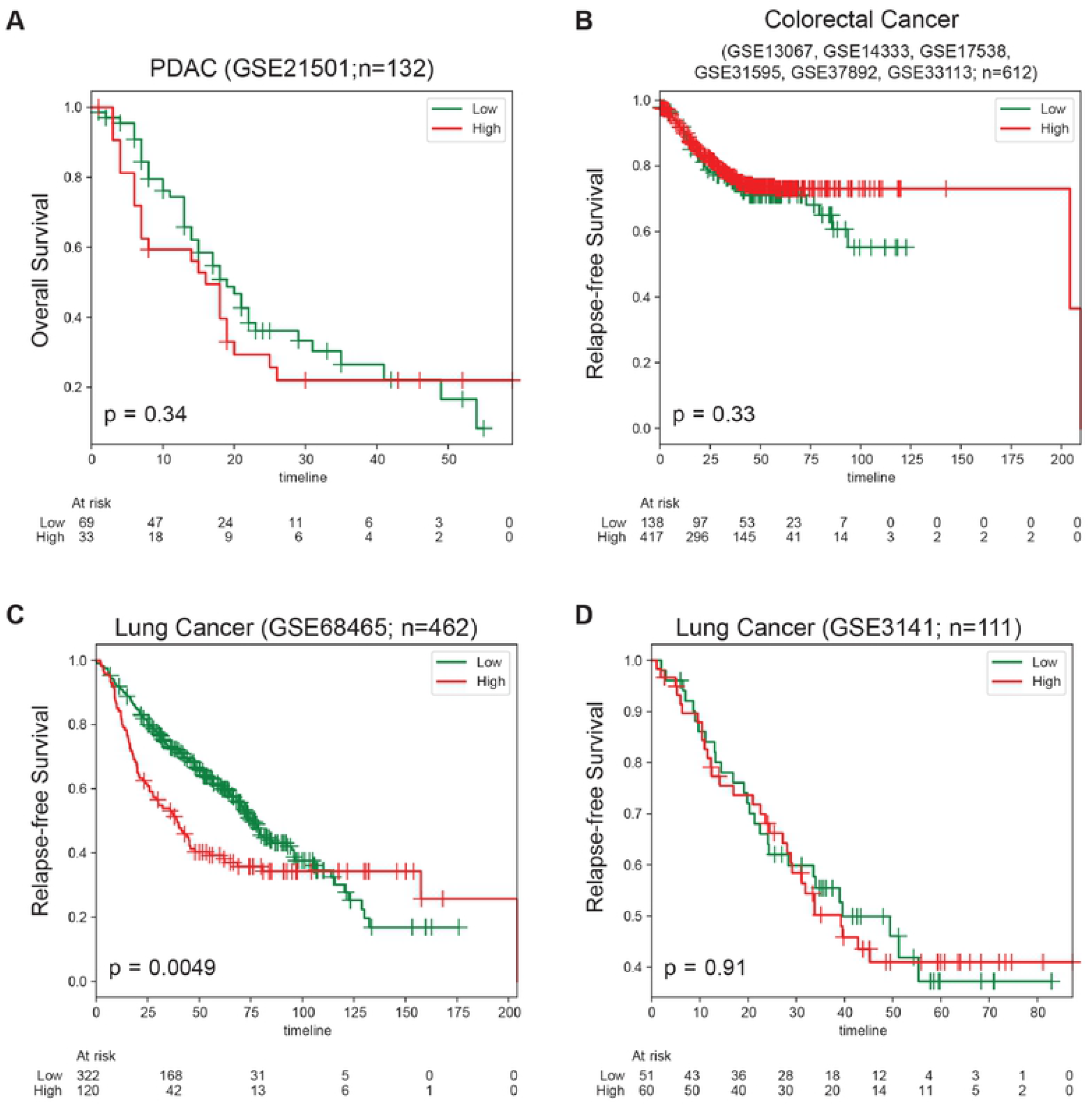
Survival outcome in diverse cancer types. iDEGs signature is used to classify cancer samples into high and low subgroups in four independent datasets (Panel A, Pancreatic Ductal Adenocarcinoma, GSE21501, n = 132; Panel B, Colorectal Cancer, n= 612; Panel C, Lung Cancer, GSE68465, n = 462; Panel D, Lung Cancer, GSE3141, n=111). (A-D) Kaplan-Meier analysis in performed using python lifelines package and p values are computed using log-rank test. Both analyses are independently verified using R statistical software.

